# Bioinformatic Mining of Novel Lipopeptides Enabled by Dap-tomycin Cs Domain and Structural Modeling

**DOI:** 10.1101/2025.06.21.660876

**Authors:** Wei Bai, Hao Zhu, Mei L. Ding, Ning Chen, Dan Y. Zhu, Nan X. Wang, Ren X. Tan

**Affiliations:** Synthetic Biology Center for Chinese Medicine, School of Pharmacy, Nanjing University of Chinese Medicine, Nanjing 210023, P. R. China; State Key Laboratory of Pharmaceutical Biotechnology, Institute of Functional Biomolecules, Nanjing University, Nanjing 210023, P. R. China

**Keywords:** lipopeptides, bioinformatic platform, Cs domain, sequence similarity network

## Abstract

Amidst the escalating crisis of antibiotic resistance, lipopeptides have emerged as promising therapeutic candidates due to their unique amphipathic structures. In this study, we developed a systematic bioinformatic platform using the condensation starter (Cs) domain of daptomycin as a molecular probe. Sequence similarity network (SSN) analysis identified 613 potential lipopeptide biosynthetic gene clusters (BGCs), with 432 (70.5%) originating from *Streptomyces* species. Subsequent integration of antiSMASH boundary prediction and evolutionary analysis prioritized 33 candidate BGCs harboring multiple post-modification modules. Five novel BGC types were ultimately selected based on Cs domain homology (<40% identity) and modification complexity. AlphaFold3 modeling revealed that WP_386473946.1 possess distinctive loop architecture, an expanded catalytic cavity, and a unique Asp60-Ala90 structural unit. Crucially, glycine residues adjacent to the conserved HHxxDG motif in the active pocket provide key targets for substrate recognition and rational engineering. This work delivers structure-guided genomic resources and molecular blueprints for accelerated discovery of antidrug-resistant lipopeptides.

## 1. Introduction

Antimicrobial resistance (AMR) has emerged as a critical global public health threat in the 21st century. World Health Organization (WHO) data reveal that drug-resistant bacterial infections already cause approximately 1.27 million deaths annually worldwide. This figure is projected to exceed 10 million deaths (Figure 1A) by 2050 without effective interventions, accompanied by economic losses surpassing 100 trillion US dollars[1]. The crisis stems from a dual imbalance, the stagnation of drug development and the relentless evolution of pathogens. Over the past two decades, only 33 novel antibiotics have been approved (Figure 1B). Metagenomic studies revealing that 23.78% of identified genes are multidrug-resistant genes underscore the rapid evolution and dissemination of antibiotic resistance. As the development rate of new antimicrobial agents lags far behind the adaptive evolution of pathogens[2], the development of novel anti-resistance drugs based on innovative molecular mechanisms[3] has become a strategic imperative in the biomedical field.

**Figure 1.**
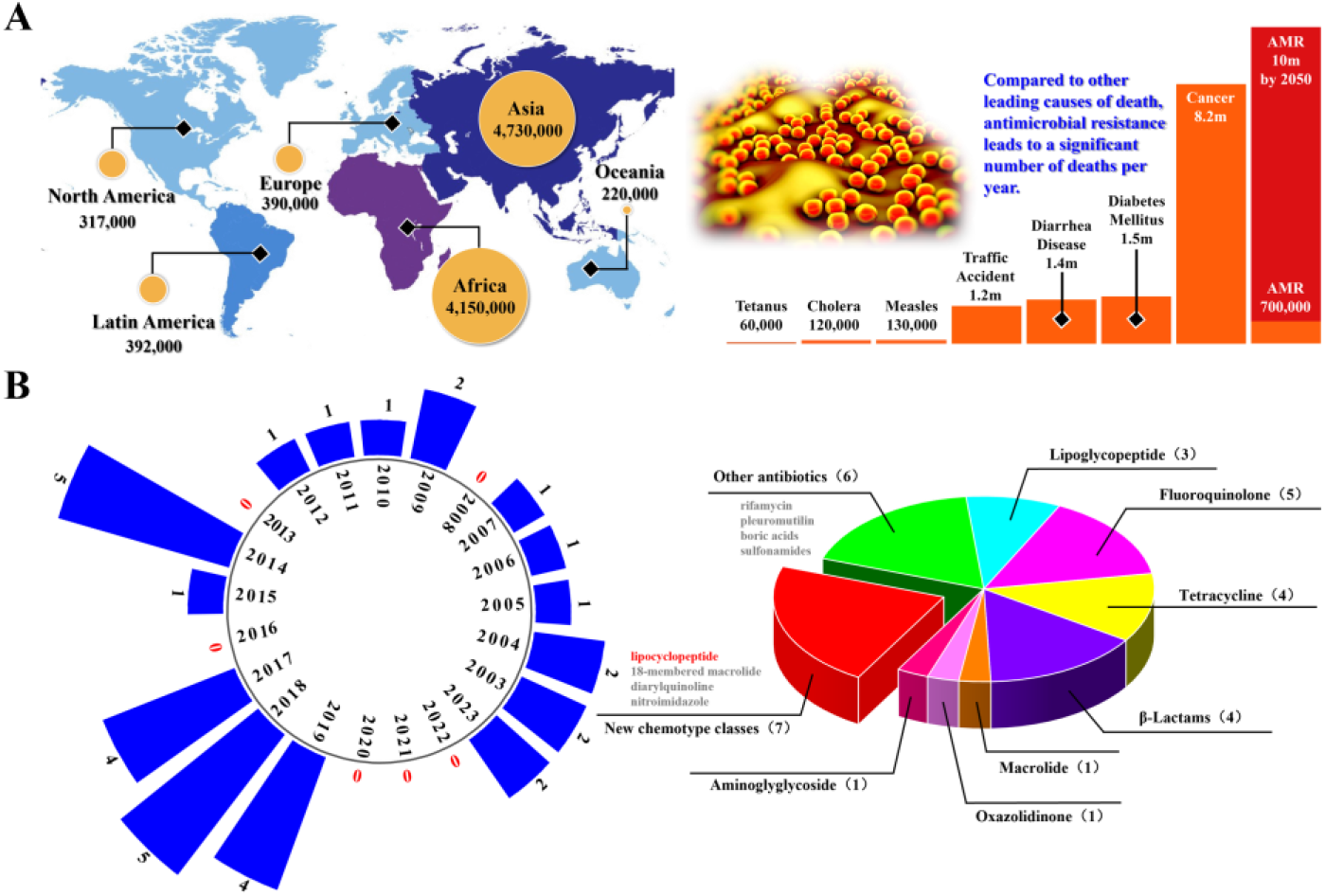
The proliferation rate of drug-resistant pathogens substantially outpaces the development rate of novel antibiotics. (A) Global incidence of multidrug-resistant infections and projections across six continents[1]; (B) FDA-approved antibiotics in the past 20 years[1].

Lipopeptide represent a promising breakthrough in combating drug-resistant bacteria due to their unique membrane-targeting mechanisms[4]. Their amphipathic architecture, featuring a peptide backbone linked via an amide bond to a fatty acid chain, enables disruption of bacterial cell membranes or inhibition of critical physiological processes[5–7]. For instance, daptomycin exerts its bactericidal effects by targeting the membrane of Gram-positive bacteria in the presence of Ca^2+^, perturbing lipid organization and inhibiting peptidoglycan biosynthesis[8,9], while globomycin competitively inhibits signal peptidase to block outer membrane synthesis towards Gram-negative pathogens[10]. Despite broad-spectrum potential of lipopeptides, daptomycin has been the only calcium-dependent antibiotic approved for clinical use in the past two decades[1], revealing the inefficiency of traditional phenotype-based screening strategies and highlighting the urgent need for structure-guided and systematic discovery platforms. This imperative necessitates comprehensive investigation into lipopeptide biosynthesis pathways. The bioassembly of lipopeptides hinges on the modular machinery of non-ribosomal peptide synthetases (NRPS) and the substrate specificity of condensation starter (Cs) domains. NRPS enzymes operate via an assembly-line process, precisely orchestrating amino acid activation (A domain), thiolation (T domain), and condensation (C domain), while relying on tailoring enzymes to incorporate non-proteinogenic units[11,12] (e.g., methylated glutamate[13]). Notably, the Cs domain acts as the gateway for fatty acid chain incorporation, catalyzing polyketides acylation through fatty acyl-AMP ligase (FAAL)[14], fatty acyl CoA ligase (FACL)[15], or polykedtide synthase (PKS)-associated pathways[11]. The architecture of the fatty acyl chain serves as the primary structural determinant governing the bioactivity of lipopeptides. Prioritizing Cs in mining pipelines therefore accelerates the discovery of novel bioactive lipopeptides. Genomic mining of uncultured soil microbes by the Sean Brady group revealed macolacin[16] and cilagicin[17] as potent anti-MRSA antibiotics, which paves a new path for combating the increasingly serious challenge of antibiotic resistance.

We employed the Cs domain of daptomycin as a molecular probe for genomic mining in this work, establishing a systematic platform for lipopeptide antibiotic discovery. Leveraging sequence similarity network (SSN) analysis centered on the daptomycin Cs domain, we identified 37 novel candidate biosynthetic gene clusters (BGCs) encoding multiple tailoring enzymes from a pool of 613 potential BGCs. Subsequent BGC evolutionary classification and AlphaFold3-based structural modeling pinpointed a novel Cs domain (WP_386473946.1) within *Streptomyces zaomyceticus*, possessing an extended loop region beyond the V-shaped catalytic pocket conformation. This integrated strategy provides a molecular blueprint for designing next-generation lipopeptides with enhanced membrane penetration and anti-resistance properties. which may serve as an innovative solution to the global AMR crisis.

## 2. Materials and Methods

### 2.1. Bibliometric Analysis

This study employed a systematic bibliometric approach to conduct a visual analysis of lipopeptide antibiotic research within the biomedical field. Literature retrieval was performed using the Web of Science Core Collection database. The search strategy was defined as: Topic sentences (TS) = (“lipopeptide” AND “antibiotic” AND “bio*”), with the publication date range restricted to January 2021 through December 2025. The retrieved records were exported in batches as plain text files (Supplementary Materials, download_1.txt to download_5.txt). Following import into CiteSpace (version 6.3.R1) and removal of duplicate entries, the time slicing parameter (“Year Per Slice”) was configured as 1-year intervals spanning from January 2021 to December 2025. Utilizing the keyword co-occurrence network analysis mode, a knowledge map was constructed with the g-index set to k = 20.

### 2.2. Construction of Sequence Similarity Network (SSN) and Cluster Analysis

BGCs of characterized lipopeptide antibiotics[18] were systematically retrieved from the MIBiG database[19] (https://mibig.secondarymetabolites.org/). Representative amino acid sequences (known.fasta) were downloaded and integrated with candidate sequences identified in this study through mining of the daptomycin Cs domain. Following redundancy removal, a non-redundant dataset of Cs sequences (Cs.fasta) was compiled. This dataset was submitted to the EFI-EST platform[20] (https://efi.igb.illinois.edu/efi-est/) for construction of a SSN using default parameters. Within the SSN, nodes represent individual protein sequences, and edge weights are defined by pairwise sequence similarity scores. The inherent clustering algorithm of the platform was employed to analyze network topology, partitioning clusters at a filter value threshold of 80% to merge low-confidence edges. Final network data (node attribute tables and edge lists) were imported into Cytoscape (v3.10.3) for visualization of key functional clusters and core hub proteins.

### 2.3. Boundary Prediction and Functional Annotation of Gene Clusters

BGC boundary prediction and functional annotation were performed using the antiSMASH (v8.0) online platform[21] (https://antismash.secondarymetabolites.org/#!/start) based on the candidate protein dataset (Supplementary Materials, id_protein_50k.fasta). BGCs containing tailoring enzymes (e.g., P450s, methyltransferases, and oxidoreductases) with high abundance of modification modules were selected. Following deduplication, a non-redundant dataset of candidate BGCs (Supplementary Materials, CS_selection.fasta) was constructed for downstream analyses.

### 2.4. Sequence Alignment and Phylogenetic Tree Construction

Multiple sequence alignment of the non-redundant candidate BGC dataset (Supplementary Materials, CS_selection.fasta) was performed using MEGA11 (v11.0) with the ClustalW algorithm to compute sequence similarity matrices. The alignment results were visualized through the ESpript 3.0 online platform[22] (https://espript.ibcp.fr/ESPript/ESPript/index.php), generating annotated alignment maps with conserved domain markers. Subsequently, the aligned sequences were subjected to maximum likelihood (ML) phylogenetic reconstruction implemented in MEGA11. The resulting tree was topologically refined and annotated using the iTOL (v6) online tool[23] (https://itol.embl.de/), with adjustments to branch length scaling and application of color-coded node labels to enhance visual interpretation of functional clusters and evolutionary relationships.

### 2.5. Prediction of Three-Dimensional Models

The three-dimensional structure of the Cs domain in target protein WP_386473946.1 was predicted using the AlphaFold3 web server[24] (https://alphafoldserver.com/welcome). Models were filtered based on per-residue plDDT confidence scores (0–100 scale), retaining regions with high-confidence plDDT values for downstream analysis. Selected models were visualized and structurally refined with default parameters in PyMOL 3.1.3, including surface rendering. To investigate the evolutionary conservation of Cs domains, the predicted model was systematically aligned with experimentally determined Cs structures from seven characterized lipopeptide antibiotics: A54145, daptomycin, enduracidin, globomycin, rakicidin, taromycin A, and telomycin. Structural superposition of Cs backbones was performed using PyMOL.

## 3. Results

### 3.1. Bibliometric Analysis of Lipopeptide Antibiotics

Based on bibliometric analysis of 903 publications retrieved from the Web of Science Core Collection database, a keyword co-occurrence network was constructed using CiteSpace (Figure 2). The network comprised 253 keywords connected by 345 links, yielding a network density of 0.0108. Analysis of high-frequency keywords (Table 1) revealed that “biological control” (frequency: 72, centrality: 0.06) and “biocontrol” (frequency: 59, centrality: 0.08) constituted the core thematic foci of lipopeptide antibiotic research over the past five years, reflecting a prevailing research emphasis on biological control applications. However, although “biosynthesis” ranked 12th among high-frequency terms with 54 occurrences, its centrality metric of zero indicates a significant knowledge gap in research on biosynthetic assembly. This research deficit stands in stark contrast to the escalating global burden of multidrug-resistant (MDR) infections and the worsening AMR crisis across six continents (Figure 1A).

**Table 1.**
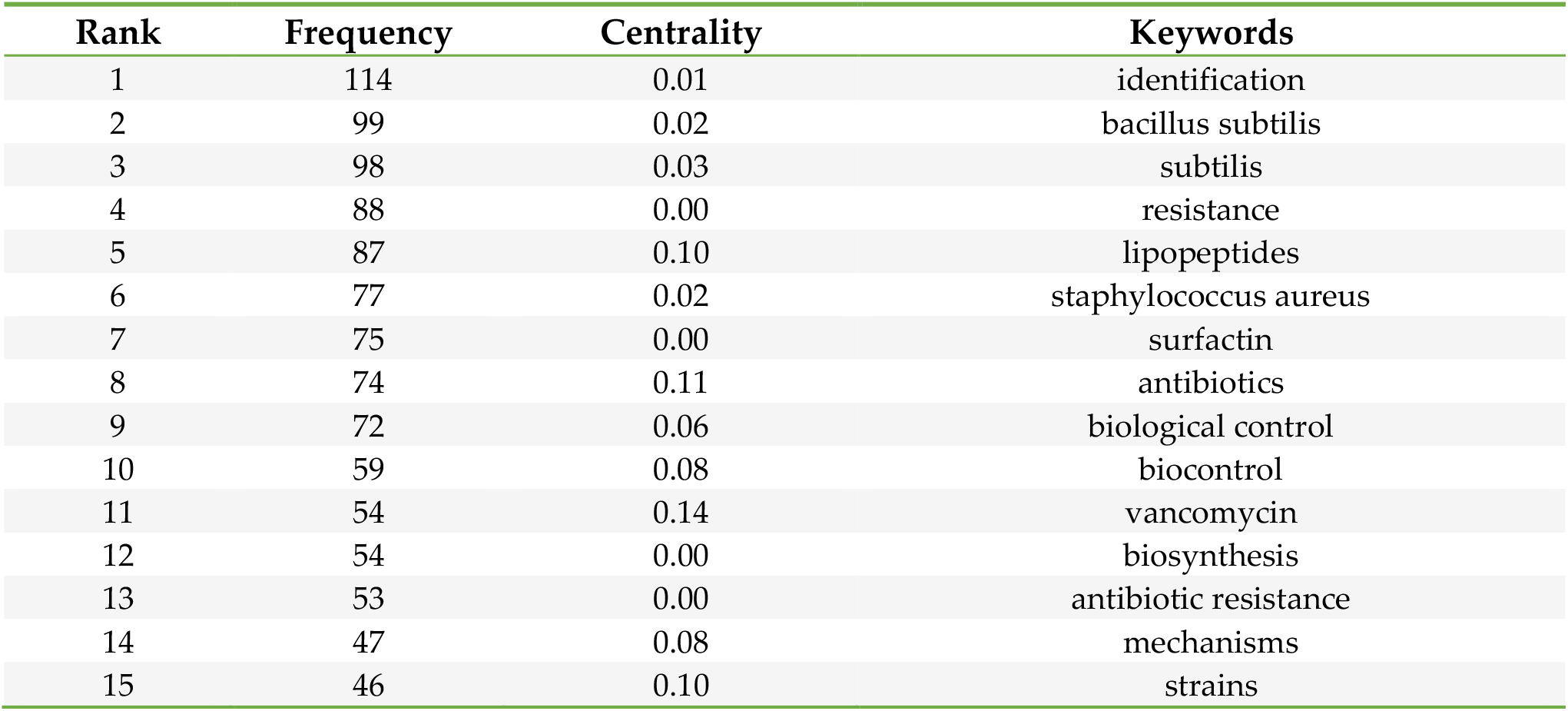
Top 15 keywords in lipopeptide antibiotic research.

**Figure 2.**
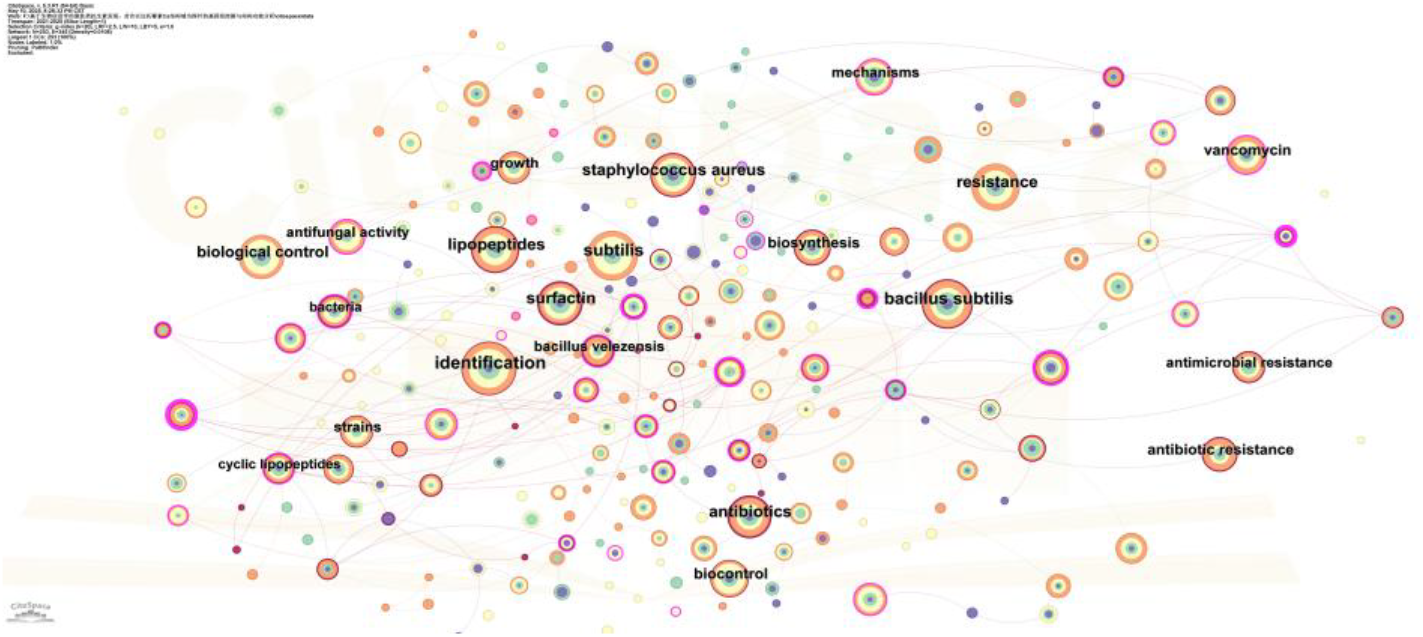
Keyword co-occurrence network for lipopeptide antibiotics research.

Critically, the pipeline for novel lipopeptide antibiotics lags far behind the accelerating pace of the rapidly evolving resistance mechanisms. Statistical data reveal that only one lipopeptide, daptomycin, was approved by the U.S. Food and Drug Administration (FDA) among 33 novel antibiotics approved over the past two decades (Figure 1B). The delayed elucidation of biosynthetic pathways and slow progress in metabolic engineering represent major bottlenecks hindering the advancement of the lipopeptide class. This developmental delay is occurring against the backdrop of a sharp increase in global incidence of AMR-related infections. Consequently, intensifying research on antibiotic lipopeptide biosynthesis is not merely pivotal for overcoming technical barriers, but an urgent imperative to address the global public health crisis. The antimicrobial potency of lipopeptide antibiotics is primarily governed by the structural architecture of their lipid tails[25]. As the Cs domain catalyzes the pivotal incorporation of these lipid moieties, targeted mining of Cs domain homologs constitutes a rational strategy for the accelerated discovery of novel bioactive lipopeptides.

### 3.2. Screening of Putative Lipopeptide BGCs

Using the Cs domain of daptomycin (Figure 3A) as a bioinformatic probe, we retrieved (>)5,000 homologous sequences from the NCBI non-redundant protein database through systematic homology searches. SSN analysis enabled clustering and redundancy reduction of these sequences (Figure 3B), ultimately identifying 613 putative lipopeptide BGCs. Strikingly, 432 BGCs (70.5% of total candidates) (Figure 3B, Supplmentary Materials) originated from *Streptomyces* species, indicating this genus dominates lipopeptide biosynthetic capacity. These results establish a valuable genetic repository for mining novel lipopeptide natural products and elucidating their biosynthetic mechanisms.

**Figure 3.**
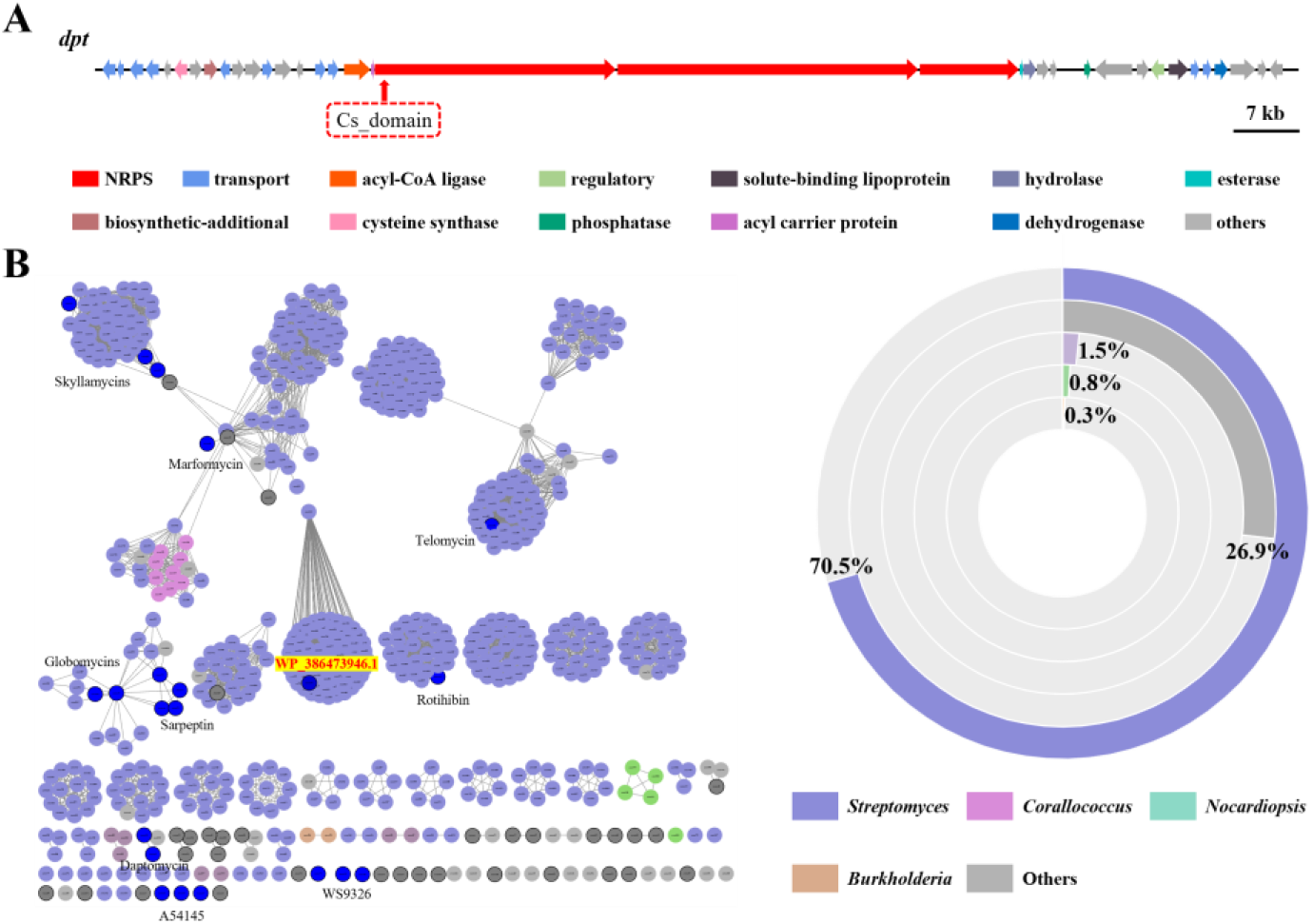
Genomic mining using daptomycin’s Cs domain as a probe within the NCBI database. (A) The BGC of daptomycin named *dpt*; (B) SSN of daptomycin Cs and its 612 homologues.

### 3.3. Evolutionary Classification of Candidate BGCs

Boundary prediction and functional annotation of target gene clusters via the antiSMASH platform enabled selection of 37 candidate BGCs harboring multiple post-assembly modification modules. Integration with 26 characterized Cs domains facilitated phylogenetic tree construction through multiple sequence alignment (Figure 4A), revealing evolutionary relationships and potential functional divergence between candidate and characterized BGCs. Application of dual screening criteria of Cs sequence identity (<40%) and complexity of modification modules prioritized five distinct BGC categories exhibiting unique structural features and high diversification potential (Figure 4B). The low sequence homology (<40% identity) strongly suggests these clusters encode novel lipopeptide scaffolds, while their complex modification machinery indicates capacity to produce structurally diverse lipopeptides with unprecedented bioactivities. These BGCs may constitute high-value targets for heterologous expression and metabolic mining.

**Figure 4.**
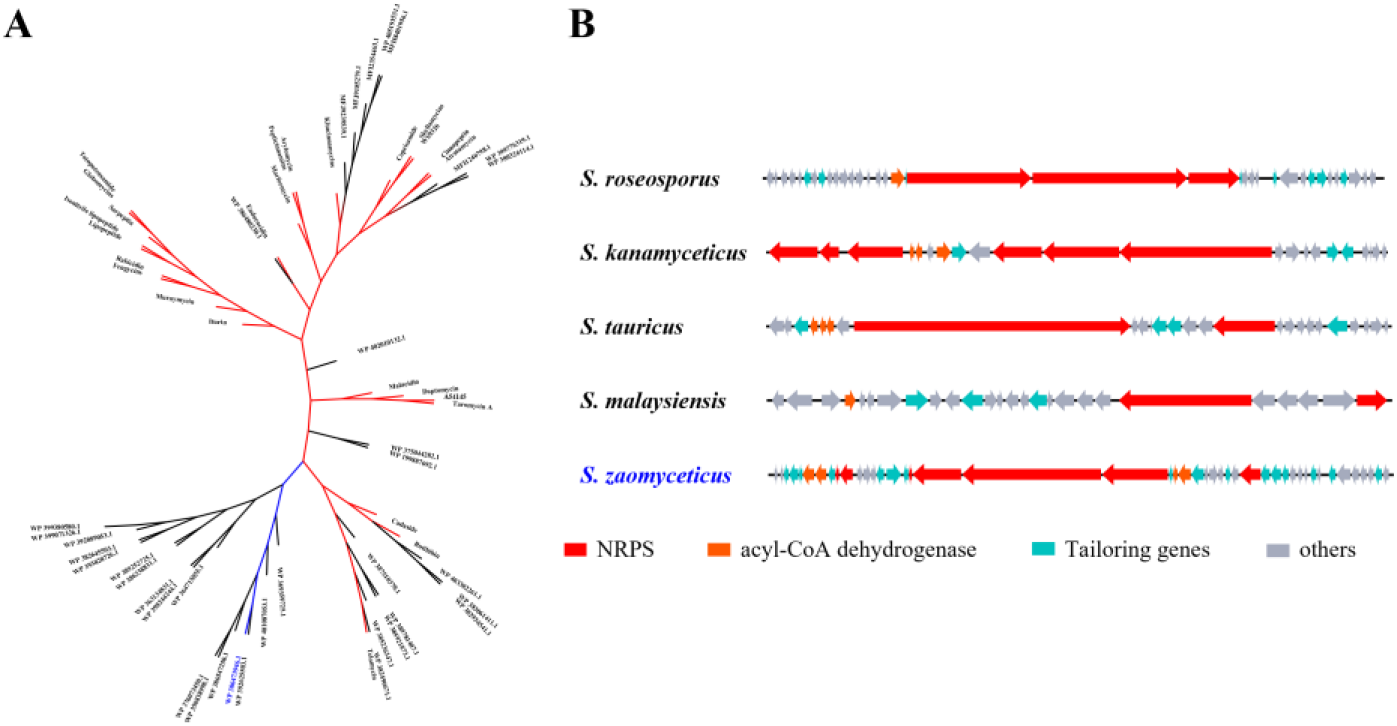
Phylogenetic analysis of distinct Cs domains. (A) Phylogeny tree of 63 Cs domains. Red branches represent characterized Cs (n = 26), black branches denote uncharacterized Cs with potential novel functions (n = 36), and blue branche corresponds to WP_386473946.1; (B) Five phylogenetically discrete BGCs classes, each demonstrating signature structural architectures and significant diversification capacity. *S. zaomyceticus* possesses WP_386473946.1.

### 3.4. Structure-Guided Functional Analysis of the Cs Domain

Phylogenetic analysis first indicated that WP_386473946.1 forms a distinct clade, suggesting it might possess a unique structure. This hypothesis was then confirmed by sequence alignment, which revealed an extended sequence segment in this Cs domain. Based on these findings, we utilized AlphaFold3 protein structure prediction platform to construct a high-confidence 3D structural model (Figure 5A). Comparative structural analysis revealed a distinct V-shaped catalytic cleft topology with significantly larger cavity volume than homologous structures. Crucially, a characteristic His59-Asp91 secondary structural element (Figure 5B) forms unique spatial arrangements likely modulating substrate-binding specificity and catalytic microenvironment. Further multiple sequence alignment identified two conserved glycine residues (Gly168/Gly170) precisely positioned adjacent to the HHxxDG functional motif within the active pocket (Figure 5C). This discovery not only provides structural rationale for substrate recognition mechanisms but also establishes critical engineering targets for rational enzyme redesign.

**Figure 5.**
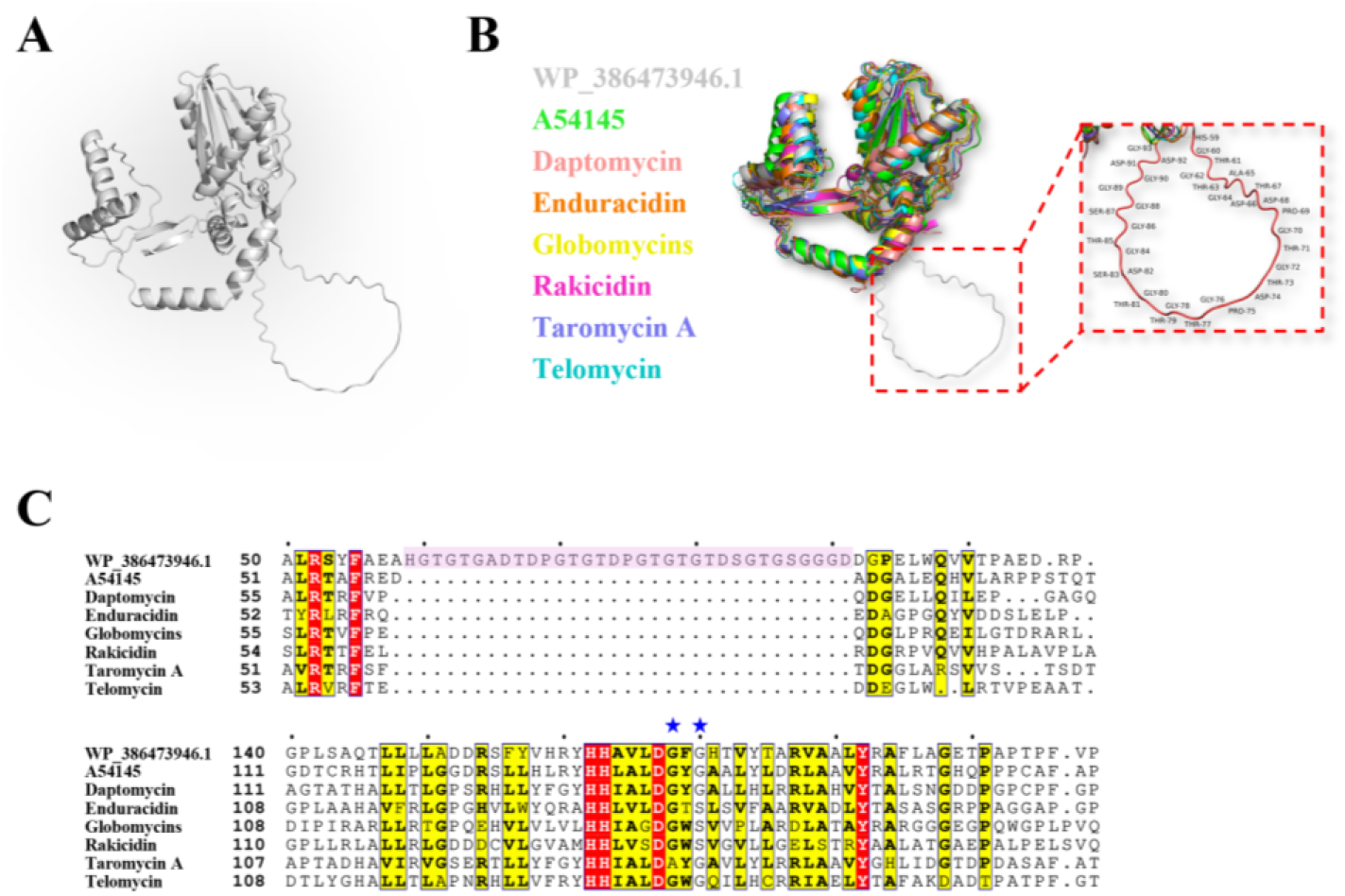
Structural prediction and sequence alignment of WP_386473946.1. (A) Three-dimensional structural modeling of WP_386473946.1 using AlphaFold3 (https://alphafoldserver.com/welcome); (B) Overlaid plot of 7 modeled known Cs and WP_386473946.1; (C) Sequence alignment of Cs of 7 known lipopeptides and WP_386473946.1.

## 4. Discussion

The escalating global crisis of AMR (Figure 1A) has intensified the demand for novel structural antibiotics with unique mechanisms of action. Lipopeptide antibiotics represent a crucial source for developing new antibiotics due to their distinctive amphipathic structure[1]. Within these molecules, the structure of the fatty acid chain introduced by the Cs domain catalysis is a key pharmacophore influencing their activity[1]. Therefore, in-depth exploration of this domain is particularly essential. This study developed a strategy for rapidly discovering novel bioactive lipopeptides. Using the daptomycin Cs domain as a probe for genome mining, we first constructed a SSN. Initial screening yielded 613 Cs domains, and cluster analysis was performed on both known and functionally uncharacterized Cs domains. The results revealed that 432 of these Cs domains originate from the genus *Streptomyces*, highlighting *Streptomyces* as a rich reservoir of lipopeptide BGCs (Figure 3B). *Streptomyces* species, owing to their inherent capacity for diverse biosynthesis driven by large genomes and rich enzymatic toolboxes such as P450 oxidases[26] and methyltransferases[27], are frequently chosen as preferred heterologous hosts. This also provides a pathway for the efficient production of structurally novel lipopeptides [28,29].

Subsequently, we conducted further analysis of the selected 432 BGCs using antiSMASH for structural prediction. This identified 37 novel lipopeptide BGCs with unknown functions and rich post-modification potential. Phylogenetic analysis was then performed comparing these 37 novel Cs domains with 26 functionally characterized Cs domains (Figure 4A). This further refined our selection to five distinct clades. These clades harbor BGCs (Figure 4B) encoding abundant post-modification enzymes and feature Cs domains exhibiting low sequence conservation, suggesting they likely encode structurally novel lipopeptides with potentially novel bioactivities awaiting for activation or heterologous expression.

Concurrently, we employed AlphaFold3 to generate three-dimensional structural models of the novel Cs domains. Notably, the model for Cs domain WP_386473946.1 revealed unique structural features (Figures 5A and 5B). A prominent extended loop region (His59-Asp91; Figure 5B) is predicted to form a specific spatial microenvironment, potentially underpinning substrate specificity. Furthermore, the co-localization of conserved glycine residues (Gly168 and Gly170) with the core HHxxDG motif within the substrate-binding pocket (Figure 5C) represents potential target sites for structural engineering[30]. Collectively, the extended loop and the conserved motifs within the binding pocket represent potential targets for the directed engineering of substrate specificity.

## 5. Conclusions

This study established a bioinformatics-driven strategy for the rapid discovery of lipopeptides, resulting in a high-value resource library of *Streptomyces* lipopeptide BGCs. From this library, we identified BGCs predicted to encode novel bioactive lipopeptides. Furthermore, three-dimensional structural predictions of candidate Cs domains directly inform engineering strategies to modulate substrate scope and selectivity. Collectively, this Cs domain-centric mining strategy provides an accelerated pathway for the discovery of lipopeptides with novel bioactivities.

## Supplementary Materials

The following supporting information can be downloaded at: https://zenodo.org/records/15669838.

## Author Contributions

Conceptualization, W. B. and H.Z.; methodology, W. B. and H.Z.; software, H.Z.; validation, M. L. D. and N. C.; formal analysis, H.Z. and D. Y. Z.; investigation, N. C. and D. Y. Z.; resources, N. X. W.; data curation, M. L. D.; writing—original draft preparation, W. B. and H. Z.; writing—review and editing, W. B. and R. X. T.; visualization, N. X. W.; supervision, R. X. T.; project administration, W.B.; funding acquisition, W. B. All authors have read and agreed to the published version of the manuscript.

## Data Availability Statement

All data generated or analyzed during this study are included in this published article and its Supplementary Materials. Portions of the data are also available in the publicly accessible databases of the NCBI (https://www.ncbi.nlm.nih.gov/).

